# Functional alterations in cortical processing of speech in glioma-infiltrated cortex

**DOI:** 10.1101/2021.05.14.444263

**Authors:** Alexander A. Aabedi, Benjamin Lipkin, Jasleen Kaur, Sofia Kakaizada, Sheantel Reihl, Jacob S. Young, Anthony T. Lee, Saritha Krishna, Edward F. Chang, David Brang, Shawn L. Hervey Jumper

**Affiliations:** Department of Neurological Surgery, University of California, San Francisco, CA, 94143, USA; Department of Psychology, University of Michigan, Ann Arbor, MI, 48109, USA

**Keywords:** glioma circuitry, neuron and glioma network, speech processing

## Abstract

Recent developments in the biology of malignant gliomas have demonstrated that glioma cells interact with neurons through both paracrine signaling and electrochemical synapses. Glioma-neuron interactions consequently modulate the excitability of local neuronal circuits, and it is unclear the extent to which glioma-infiltrated cortex can meaningfully participate in neural computations. For example, gliomas may result in a local disorganization of activity that impedes the transient synchronization of neural oscillations. Alternatively, glioma-infiltrated cortex may retain the ability to engage in synchronized activity, in a manner similar to normal-appearing cortex, but exhibit other altered spatiotemporal patterns of activity with subsequent impact on cognitive processing. Here, we use subdural electrocorticography to sample both normal-appearing and glioma-infiltrated cortex during speech. We find that glioma-infiltrated cortex engages in synchronous activity during task performance in a manner similar to normal-appearing cortex, but recruits a diffuse spatial network. On a temporal scale, we show that glioma-infiltrated cortex has lower capacity for information encoding when performing nuanced tasks such as speech production of monosyllabic versus polysyllabic words. As a result, temporal decoding strategies for distinguishing monosyllabic from polysyllabic words were feasible for signals arising from normal-appearing cortex, but not from glioma-infiltrated cortex. These findings inform our understanding of cognitive processing in chronic disease states and have implications for neuromodulation and prosthetics in patients with malignant gliomas.

**Significance Statement:** As gliomas proliferate, they infiltrate healthy brain tissue. Often, patients with such tumors in the language areas of the brain develop aphasia. Understanding how gliomas interact with normal neural circuits is critical for developing neuroprostheses that restore speech. Recent evidence demonstrates that glioma cells interact synaptically with neurons, and thus can modulate neural circuits. However, it is unclear the extent to which glioma-infiltrated cortex participates in cognitive processing. Using electrocorticography to record both glioma-infiltrated and normal-appearing cortex during speech, we found that glioma-infiltrated cortex is capable of coordinated neural responses, but has reduced capacity for information encoding. Instead, glioma-infiltrated cortex recruits a broader network of cortical regions during speech, which may represent a compensatory mechanism with implications for future neuroprostheses.

## Introduction

Gliomas are the most common intrinsic brain tumors and a leading cause of neurological impairment in adults. Mounting evidence suggests that malignant gliomas remodel functional neuronal networks, and that these cellular- and network-level interactions impact cognition and survival (1). The mechanistic underpinnings of glioma-neuron interactions and their influence on human cognition is incompletely understood. Recent studies have shown that gliomas not only interact with neurons through paracrine signaling, but also through *de novo*, electrophysiologically-coupled synapses. As such, glioma-infiltrated cortex maintains activity-dependent connections to disparate brain regions (2–4). These findings challenge the notion that infiltrative gliomas simply lead to the destruction and loss of function within tumor-infiltrated brain regions, raising instead the possibility that brain cancer may fundamentally *alter* the way infiltrated brain regions perform the computations required for cognition and behavior.

There are two primary means by which gliomas may alter the electrophysiology of the human cortex. On one hand, glioma invasion may lead to locally disorganized activity that impedes the transient synchronization of neural oscillations required for the binding of task-relevant information (5). If this were the case, glioma-infiltrated cortex would not meaningfully participate in neural computations. Alternatively, glioma-infiltrated cortex may retain the ability to engage in synchronized activity, in a manner similar to normal-appearing cortex, but suffer from a degradation in the amount of information encoded on a temporal scale. If so, a more diffuse spatial area of glioma-infiltrated cortex may be recruited to perform computations (6).

The extent to which information is processed within tumor infiltrated cortex in the human brain is unknown. A better understanding of the biology of speech within glioma-infiltrated cortex will add to our understanding of language processing and facilitate the development of novel electrophysiologically-based interventions (i.e., neuroprosthetics and neuromodulation) for the restoration and rehabilitation of neurological function (7). Here we test the hypothesis that glioma-infiltrated cortex retains event-related neuronal synchrony with an altered capacity for information encoding compared to normal-appearing cortex. To do so, we acquired invasive electrophysiologic recordings (electrocorticography; ECoG) in non-aphasic study participants with dominant hemisphere perisylvian malignant gliomas during speech production; cortical sites were spatially matched for normal-appearing and glioma-infiltrated regions. We then applied an information theoretical framework to directly compare the encoding capacity and decodability of signals arising from these regions (8).

## Results

Our goal was to examine the computational capacity and decodability of neural signals arising from normal-appearing and glioma-infiltrated cortex during speech production from non-aphasic human study participants. As such, we recorded ECoG activity in twelve non-aphasic patients with cortically-projecting malignant gliomas in perisylvian language regions during a visual confrontation naming task (**Fig. 1**). In each participant, subdural ECoG grids were implanted over the left perisylvian cortex. We then used a combination of stereotactic neuro-navigation and offline electrode registration methods to identify electrodes overlying regions of apparently normal parenchyma (i.e., devoid of T2-FLAIR signal abnormality; “normal-appearing”) and regions with glioma invasion (i.e., within T2-FLAIR signal abnormality but outside the tumor core; “glioma-infiltrated”) (**Fig. S1**) (9).

**Figure 1:**
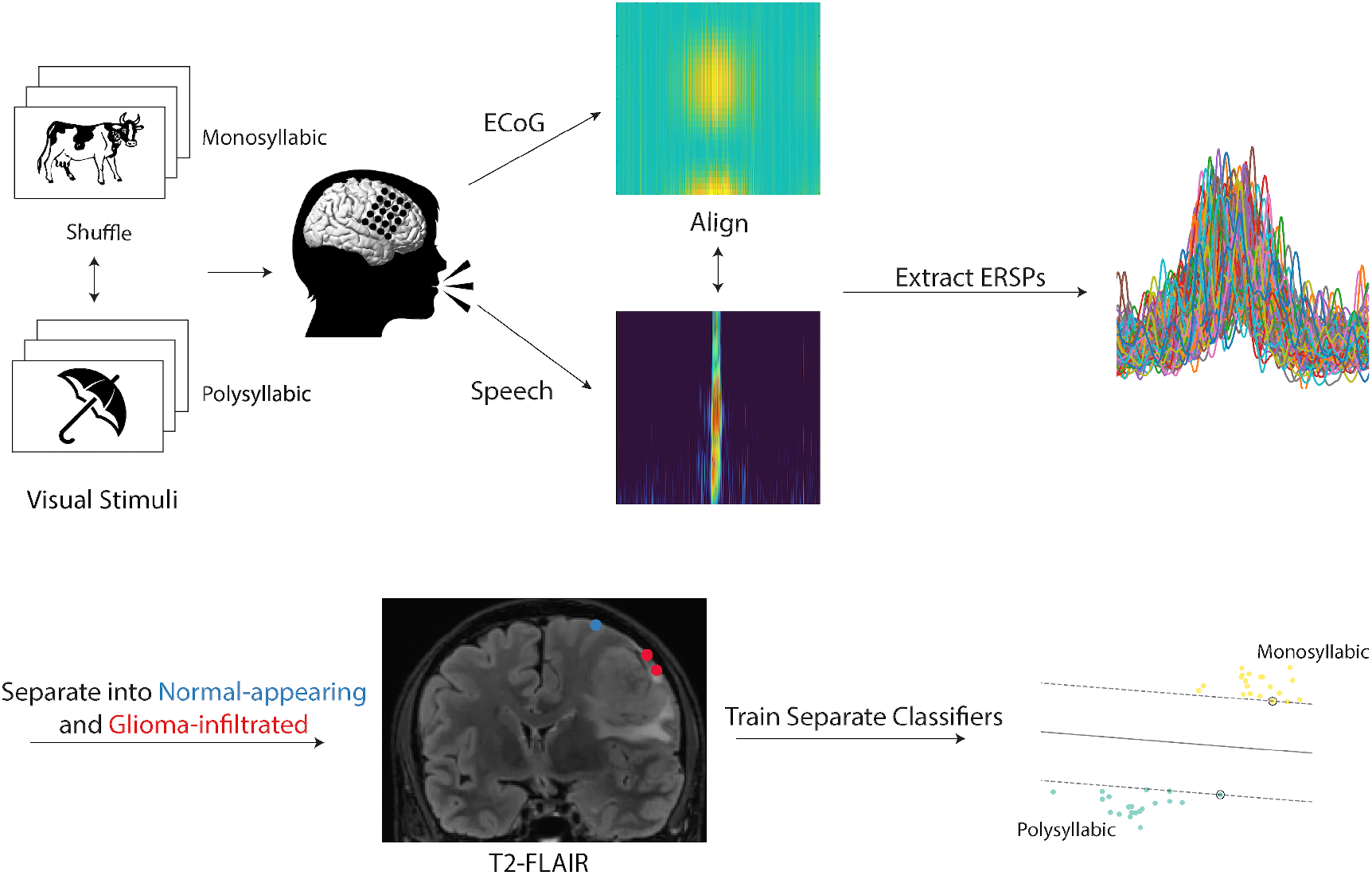
Experimental workflow. Participants with cortically-projecting perisylvian gliomas were asked to complete a picture naming task during intraoperative functional mapping. On each trial, participants were presented with a line drawing of a common object or animal in a randomized fashion and prompted to provide its name. Of the 48 stimuli, 28 represented monosyllabic words while 20 represented polysyllabic words. Audio and electrophysiologic (via electrocorticography; ECoG) recordings were taken from each participant during task completion and aligned to speech onset. Event-related spectral perturbations (ERSPs) were extracted in the high-gamma range (70 – 170 Hz). Electrodes were localized on each participant’s preoperative T2-FLAIR image and categorized as “normal-appearing” if overlying normal-appearing cortex or “glioma-infiltrated” if overlying regions of T2-FLAIR signal abnormality. Using the ERSPs, separate classifiers were trained to decode the trial condition (monosyllabic versus polysyllabic) in normal-appearing and tumor-infiltrated cortex.

Accordingly, we were able to sample from spatially matched regions and obtain excellent coverage of areas involved in the planning and articulation of speech (left inferior frontal gyrus and lateral pre-central gyrus, respectively) (10) (**Fig. 2*A-D***).

**Figure 2:**
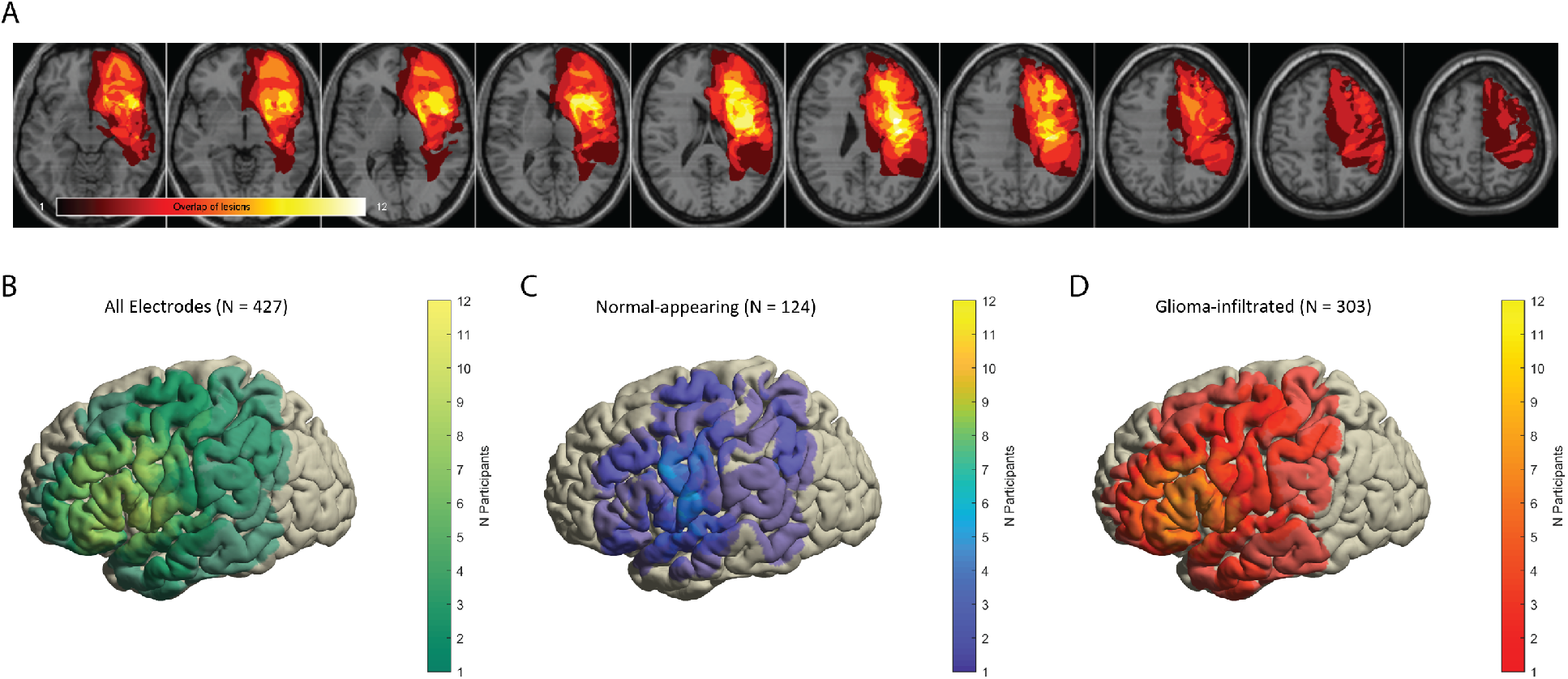
Lesion and electrode overlap. (A) Lesion overlap map using the borders of T2-FLAIR signal abnormality demonstrating involvement of the left perisylvian language areas. (B-D) common cortical surfaces showing the number of participants with electrode coverage at a given region: all electrodes (B), electrodes overlying normal-appearing cortex (C), and electrodes overlying glioma-infiltrated cortex (D). Speech planning and initiation sites are represented in both normal-appearing and glioma-infiltrated cortex.

### Glioma-infiltrated cortex engages in coordinated neural responses during speech

We extracted event-related spectral power perturbations (ERSPs) in the high-gamma range (70 – 170 Hz). During the production of speech, we found robust and coordinated responses in both normal-appearing and glioma-infiltrated cortex (**Fig. 3*A***). This includes activation anterior to primary motor cortex beginning 800 milliseconds prior to speech onset with maximal activity at the beginning of speech production (time = 0) (11). We then projected these responses onto a common cortical surface separately for normal-appearing and glioma-infiltrated cortex to examine the spatial involvement of the lateral prefrontal areas (**Fig. 3*B***). In normal-appearing cortex, we observed maximal high-gamma activity in a spatially restricted region involving the canonical speech planning areas (pars orbitalis and triangularis of the inferior frontal gyrus) (12). By contrast, in glioma-infiltrated cortex, the spatial pattern of maximal high-gamma activity included not only the rostral inferior frontal gyrus, but also diffuse areas of the middle frontal gyrus as well. These data suggest physiologically intact temporal patterns of neural responses in glioma-infiltrated cortex with broad spatial representations.

**Figure 3:**
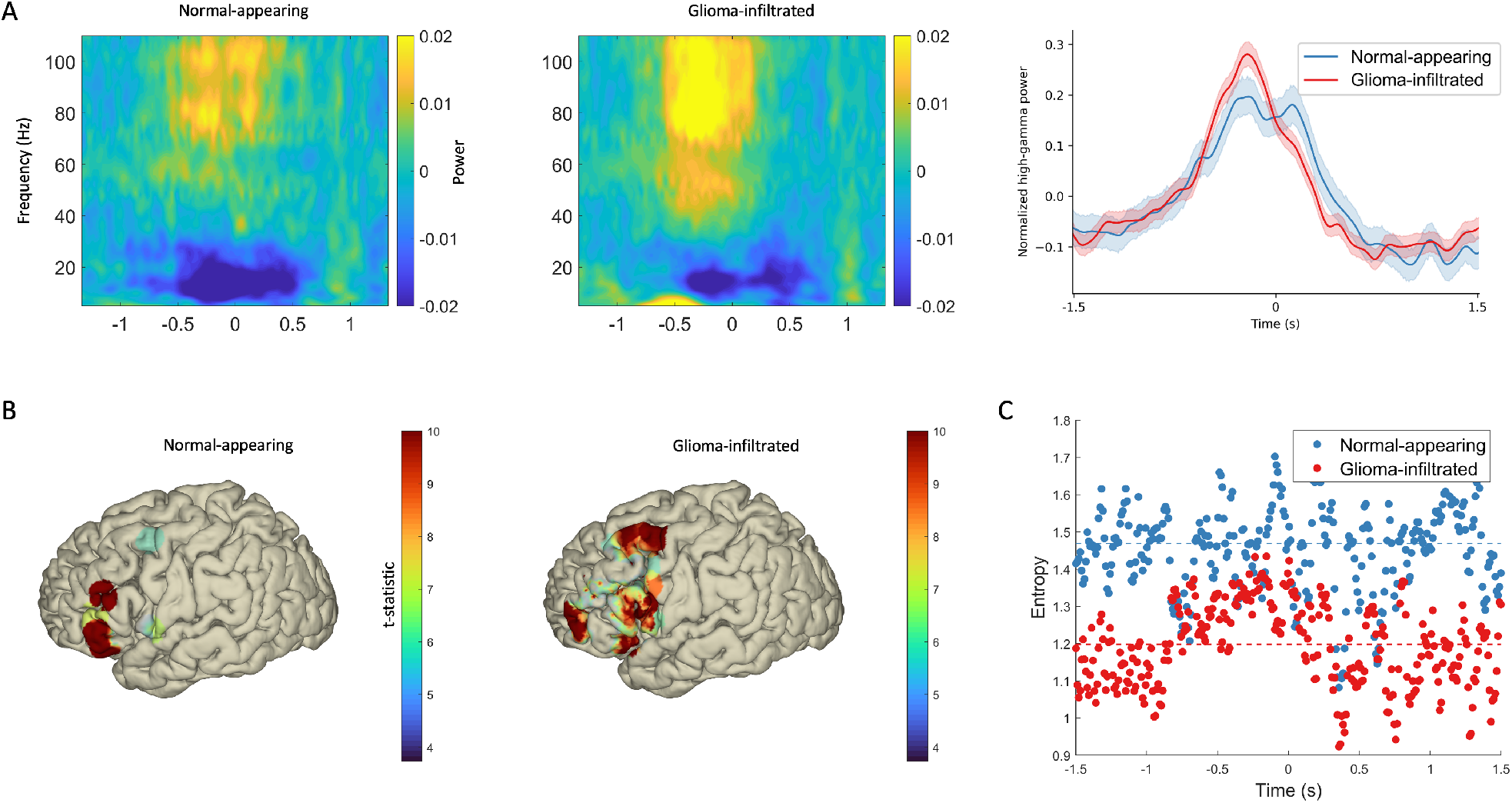
Spatiotemporal features of neural signals from normal-appearing and glioma-infiltrated cortex during speech. (A) Broadband event-related spectral perturbations (ERSPs) in normal-appearing and glioma-infiltrated cortex (left and middle, respectively). (Right) ERSPs averaged over the high-gamma range with time = 0 representing speech onset (shading represents 95% confidence intervals). Both normal-appearing and glioma-infiltrated cortex demonstrate robust and coordinated increases in time-locked high-gamma activity. (B) Spatial representations of mean high gamma activity during pre-speech planning (−1,000 to 0 ms) relative to a post-speech baseline thresholded to a Bonferroni-corrected p < 0.01. In normal-appearing cortex (left), activity is primarily restricted to the canonical speech planning sites, while activity is more diffuse in glioma-infiltrated cortex (right). (C) The encoding capacity at each time point in normal-appearing and glioma-infiltrated cortex. Signals from glioma-infiltrated cortex have lower entropy and thus less capacity to encode information (p = 0.006). Dashed lines represent sample medians.

### Glioma-infiltrated cortex has a diminished capacity for information encoding

Having established that glioma-infiltrated cortex retains the ability to engage in coordinated neural responses and recruits a diffuse network of lateral prefrontal areas during speech production, we next set out to determine the relative capacity of these cortical regions to encode information. Therefore, in this setting, information content was represented as mean high gamma activity over time within glioma-infiltrated and normal-appearing cortex. Shannon’s metric for information (Shannon entropy) provided a theoretical maximum for the signal’s encoding capacity (13). We measured the Shannon entropy of neural signals at a) each time point in the event-related response to generate a time-course of entropy values (**Fig. 3*C***) as well as b) averaged across all trials to derive a mean estimate (**Fig. S2**). Using linear mixed-effects modeling to account for the hierarchical nature of the data, we found that glioma-infiltrated cortex had lower entropy and therefore less capacity to encode information over time (p = 0.006).

### Neural signals from normal-appearing but not glioma-infiltrated cortex can be decoded

Given the coordinated neural responses, yet diminished encoding capacity of glioma-infiltrated cortex, we next set out to determine whether there are alterations in the high gamma activity elicited by computationally demanding tasks. Vocalization of polysyllabic words, for instance, requires a more intricate coordination of articulatory elements than that of monosyllabic words. The reliability of sub-lexical articulatory encoding within sensorimotor cortex has been well established (14). We therefore postulated that in normal-appearing cortex, differences in the representations of monosyllabic and polysyllabic words would be apparent in the neural signals restricted to the lateral primary motor area (M1). We separated trials in which participants vocalized monosyllabic words from those in which they vocalized polysyllabic words (**Fig. 4*A***). In normal-appearing cortex, we found greater mean high-gamma activity during polysyllabic trials (460 – 520 ms, p < 0.05; Bonferroni-corrected) when compared to monosyllabic trials. Linear mixed-effects modeling confirmed that mean high-gamma activity during vocalization of polysyllabic words was significantly higher than that during vocalization of monosyllabic words. However in glioma-infiltrated cortex, differences between polysyllabic and monosyllabic trial conditions were not apparent.

**Figure 4:**
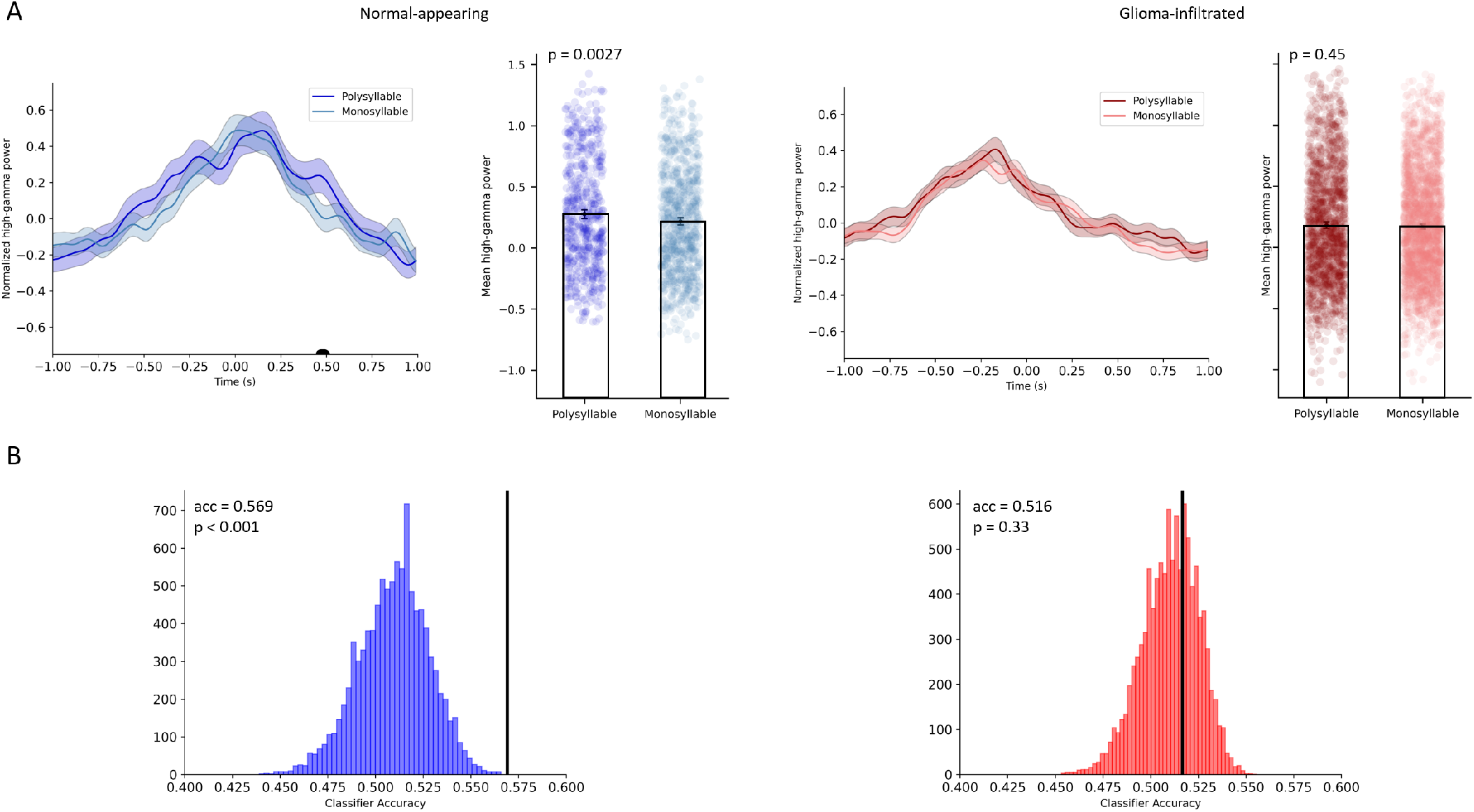
Decodability of signals from normal-appearing and glioma-infiltrated cortex in motor areas. (A) In normal-appearing cortex, ERSPs during vocalization of polysyllabic words demonstrate unique temporal features. Using linear mixed-effects modeling, mean high gamma activity in normal-appearing cortex was found to be significantly higher during vocalization of polysyllabic words (p = 0.0027). In glioma-infiltrated cortex, no statistically significant differences in high-gamma activity were found between polysyllabic and monosyllabic trial conditions. Black bar indicates time points in which the t-test p-value exceeded a threshold of p < 0.05 after Bonferroni corrections. (B) Above-chance decoding of monosyllabic from polysyllabic trials was achieved across in normal-appearing cortex (left) but not in glioma-infiltrated cortex (right) using a Bernoulli Naïve Bayes classifier. Decoding accuracy (acc; black bars) was significantly higher in normal-appearing cortex compared to glioma-infiltrated cortex (p = 0.019).

We confirmed this finding by evaluating the decodability of signals arising from normal-appearing and glioma-infiltrated cortex. We trained a Bernoulli Naïve Bayes classifier to distinguish between monosyllabic and polysyllabic trial conditions using event-related responses (**Fig. 4*B***). We implemented identical training and cross-validation paradigms (i.e., leave-one-subject-out-cross-validation) for both conditions. As expected, normal-appearing cortex produced above-chance decoding between monosyllabic and polysyllabic trials (mean classifier accuracy = 0.569, p < 0.001). By contrast, in glioma-infiltrated cortex, we were not able to decode monosyllabic from polysyllabic trials above chance (mean classifier accuracy = 0.516, p < 0.33). This pattern was maintained for support vector machine and artificial neural network decoders (**Fig. S3**). Linear mixed-effects modeling confirmed that decoding accuracies were significantly higher in normal-appearing cortex compared to glioma-infiltrated cortex (p = 0.019).

## Discussion

Emerging developments into the cellular- and network-level electrophysiologic interactions between neurons and gliomas raise critical questions about how altered neuronal circuits represent and encode information in humans. Gliomas form interconnected networks (15) that substantially modulate the excitability of neurons (9) through paracrine signaling and synaptogenesis (3, 16). Therefore, the consequences of glioma infiltration for neuronal processing could range anywhere from the loss of event-related synchronicity to the relative preservation of circuit dynamics. Here, we show that glioma-infiltrated cortex retains the ability to engage in coordinated neural responses during the planning and production of speech, but has a diminished capacity for information encoding which may stem from glioma synaptic signaling that biases the activity of these networks at baseline (9). As a result, salient features of a given cognitive/behavioral event (in this case, vocalization of mono-versus polysyllabic words) could not be temporally decoded from its neural signals. Glioma-infiltrated cortex may alternatively rely on more spatially distributed encoding to represent speech information. This prompts an exploration of how neuron-glioma communication contributes to reduced functional states and highlights the need for more targeted analysis of neuromodulation in patients with malignant gliomas.

We arrived at these findings by first demonstrating via subdural ECoG that glioma-infiltrated cortex is capable of synchronizing its activity to the onset of speech. This is in line with evidence from the neurosurgical literature where functional cortex in regions of glioma infiltration has been described during brain mapping (17, 18). For instance, evoked magnetic responses have been shown in glioma-infiltrated sensorimotor cortex after tactile stimulation (19); motor and language function can be transiently inhibited and localized via direct electrical stimulation of glioma-infiltrated cortex (20, 21); and resection of glioma-infiltrated regions with high functional connectivity to the rest of the brain leads to permanent neurological injury (22, 23). However, the spatiotemporal dynamics underlying these phenomena have, up until now, remained largely unexplored. These findings directly demonstrate that glioma-infiltrated cortex participates in coordinated neural responses during speech production while maintaining that its spatial and temporal patterns of activity are indeed distinct from normal-appearing cortex.

This is especially apparent from the comparative spatial distributions of high-gamma activity in normal-appearing and glioma-infiltrated cortex. Under physiologic conditions, pre-speech activity during visual confrontation naming localizes to a well-defined circuit, including the pars orbitalis and pars triangularis of the inferior frontal gyrus where semantic-to-lexical mapping takes place (24, 25). Here, we show that normal-appearing cortex roughly adheres to this canonical network as the regions of maximal high-gamma activity were centered on the rostral inferior frontal gyrus. Conversely, we found that glioma-infiltrated cortex recruited a diffuse network of atypical cortical regions during speech planning, including caudal portions of the middle frontal gyrus. Parallel reports of cortical redistribution of language function to atypical peritumoral sites, including the middle frontal gyrus, have been reported with PET (26, 27) and fMRI (28), but chiefly attributed the phenomenon to physiologic neuroplasticity (18, 29). In the context of the present study and evidence of coordinated high-gamma responses in glioma-infiltrated cortex, the widespread recruitment of perisylvian cortex in such regions may reflect activation of the inherently diffuse, interconnected neuron-to-glioma network (30). Further, this increased spatial representation may act as a potential short-range mechanism of native compensation for the loss of temporally encoded information in glioma-infiltrated cortex.

The next critical question in light of this evidence is, if glioma-infiltrated cortex can generate robust synchronous activity during speech in a manner similar to normal-appearing cortex, why do patients with infiltrative gliomas often present with aphasia (i.e., speech impairment)? Prior accounts of the pathophysiology of aphasia in patients with glioma have mainly focused on disruptions in anatomic and functional connectivity involving the perisylvian language network (31, 32). Here we provide a more mechanistic account by demonstrating that the neuronal subpopulations within glioma-infiltrated cortex have inherently lower capacities for information encoding given their diminished response variability/entropy (**Fig. 3*A-B***) which may stem from glioma synaptic signaling that biases the activity of these networks at baseline (9). Indeed, the diminished encoding capacity of glioma-infiltrated cortex, even in non-aphasic patients, may be indicative of alterations in cognitive processing, that when further deteriorated, may lead to aphasia. This is particularly relevant in the context of neuromodulation and neuroprostheses, as many non-aphasic patients with glioma will become aphasic over the course of their disease trajectory.

By providing a model for network-level alterations in glioma-infiltrated cortex, these findings offer insight into a potential mechanism of native compensation in non-aphasic patients. Currently, treatment options for patients with aphasia include compensatory training, environmental modification interventions, as well as assistive devices which rely on residual oral facial movements. Brain-computer interfaces directly record cortical brain signals to replace lost function and represent a new generation of rehabilitative technologies. However, the application of neuroprostheses may not be generally applicable across diseases, as loss of speech can occur in both acute and chronic neurological conditions (e.g., gliomas, brain metastases, traumatic brain injury, stroke, and primary progressive aphasia). A mechanistic understanding of disease-specific speech biology as well as associated compensation strategies is of critical importance. For glioma in particular, the increasing number of long-term survivors makes understanding the underlying effects of gliomas on network dynamics a foundational area of investigation for future therapies.

## Materials and Methods

### Experimental Workflow

To assess the computational capacity of glioma-infiltrated cortex with respect to language, invasive electrophysiologic recordings were taken from twelve adult patients with cortically-projecting perisylvian gliomas while they engaged in a visual confrontation naming task during awake craniotomy. On each trial of the task, participants were presented with a line drawing of a common object or animal and were prompted to vocalize its name. Each participant received a total of 48 unique trials in a single-block design and the correct speech responses for the trials were matched based on whether they were monosyllabic (N = 28) or polysyllabic (N = 20). Trials were randomized for each participant and manually advanced by a clinical research coordinator after a response was given. Each participant received a training session two days prior to participation to ensure familiarity with the task and underwent 1) a minimum anesthesia washout period of twenty minutes and 2) an extensive post-emergence wakefulness assessment to ensure adequate arousal during intraoperative language testing (33). Tasks were delivered via PsychToolbox 3 (http://psychtoolbox.org/) in the operating room on a 15-inch monitor (60 Hz refresh rate) placed 30 cm from the participant. All participants provided written informed consent to participate in this study, which was approved by our institutional review board (CHR-17-23215) and performed in accordance with the Declaration of Helsinki.

### Signal Acquisition and Pre-processing

To extract local cortical responses during a speech event of interest (i.e., vocalization of a monosyllabic word), participants underwent simultaneous audio and electrophysiologic recordings. Audio was sampled at 44.1 kHz from a dual-channel microphone placed 5 cm from the participant and electrophysiologic signals were amplified (g.tec; Schiedlberg, Austria) and sampled at 4800 Hz from multichannel low-density (20 electrodes, 10 mm spacing) and high-density (96 electrodes, 3 mm spacing) platinum subdural electrocorticography grids (Ad-Tech; Oak Creek, Wisconsin). During offline analyses, audio and electrophysiologic recordings were manually aligned, resampled, and segmented into epochs (speech-locked). These epochs set time = 0 ms as speech onset and included ±2,000 ms for a total of 4,000 ms of signal per trial. Trials were discarded if a) an incorrect response was given (including fillers and interjections) or b) there was a >2 second delay between stimulus presentation and response so as to maintain consistent trial dynamics and ensure that the neural signal indeed reflected the experimental manipulations. Noisy channels were automatically rejected if their kurtosis exceeded 5.0. Electrodes were subsequently re-referenced to the common average for each participant.

Because transient increases in high-gamma power are strongly correlated with the spiking activity of local ensembles and thus reflect fundamental units of neural computation (34, 35), we focused our subsequent analyses on event-related spectral perturbations (ERSPs) in the high-gamma range. To extract the ERSPs, electrophysiologic signals were first down-sampled to 600 Hz, then high-pass filtered at 0.1 Hz to remove DC-offset and low frequency drift, notch filtered at 60 Hz and its harmonics to remove line noise, and bandpass filtered between 70 and 170 Hz (i.e., the high-gamma range) using a Hamming windowed sinc FIR filter. Spectral power was extracted as the squared magnitude of the analytic signal, calculated using the Hilbert transform. These signals were finally smoothed using a 100ms Gaussian kernel, down-sampled to 100 Hz, and z-scored across each trial. All signal pre-processing was conducted using EEGLAB(36) (version 2021.0; https://sccn.ucsd.edu/eeglab/).

### Grid Localization and Identification of Glioma-infiltrated Cortex

To determine whether a given electrophysiologic signal originated from glioma-infiltrated or normal-appearing cortex, the subdural electrocorticography grids were localized via a combination of intraoperative photography and stereotactic techniques (37, 38). For each participant, intraoperative photographs of the cortex taken with and without the grid(s) were used to register the grids to reconstructions of the pial surface and vascular anatomy and extract the three-dimensional coordinate of each electrode. Each electrode coordinate was then registered to the T2-weighted FLAIR image and labeled as “glioma-infiltrated” if found overlying a region of peritumoral FLAIR signal abnormality, or “normal-appearing” if found overlying normal parenchyma by a trained co-author blinded to the electrophysiologic data (9). These labels were reviewed by a board-certified neurosurgeon (SLH-J) and compared to labels derived during intraoperative stereotactic neuro-navigation to reach a consensus (Brainlab; Munich, Germany). Finally, electrode coordinates were registered to a common surface (MNI template) for each participant to facilitate group comparisons and regions of interest were defined according to the Automated Anatomical Labeling atlas (39) (https://www.gin.cnrs.fr/en/tools/aal/). Image registration and normalization routines were conducted using SPM12 (https://www.fil.ion.ucl.ac.uk/spm/software/spm12/) and ANTsR (version 0.40; https://github.com/ANTsX/ANTsR) and surface reconstruction using FreeSurfer (40) (version 7.1; https://surfer.nmr.mgh.harvard.edu/). The location of grid implantation was solely directed by clinical indications.

### Information Theoretical Analysis

Shannon entropy was used to determine the capacity of glioma-infiltrated cortex to encode task-relevant information compared to normal-appearing cortex. In general, systems with high entropy demonstrate substantial variability across trial conditions and are thus capable of encoding more information than those with low entropy (13, 41). Two methods were used to measure the entropy of signals originating from normal-appearing and glioma-infiltrated cortex. First, Shannon entropy was computed at each time point in the ERSPs simultaneously across all trial conditions to extract time courses of encoding capacity of normal-appearing and glioma-infiltrated cortex (Neuroscience Information Theory Toolbox, https://github.com/nmtimme/Neuroscience-Information-Theory-Toolbox) (42). Next, the Shannon entropy was calculated separately for each trial to derive estimates of overall response variability of normal-appearing and glioma-infiltrated cortex by discretizing each ERSP into probability mass functions consisting of 10 equally spaced bins (8) (Scipy, version 1.6.1; https://www.scipy.org/).

### Neural Decoding

Finally, the decodability of signals arising from normal-appearing and glioma-infiltrated cortex was determined by training classifiers to distinguish between monosyllabic and polysyllabic trials across all participants. For this analysis, only electrodes from motor cortex were selected, as they are most relevant to predictability of articulation. Glioma-infiltrated and normal-appearing electrodes were separated into two groups, and the following steps were repeated exactly the same for both groups. Features were composed of electrode-level time-series of individual trials, and targets were encoded as a binary label of trial-type (1 for monosyllabic, 0 for polysyllabic). Given its prior use to quantify information encoding in neural populations and its parallels to the Shannon entropy method, a Bernoulli Naïve Bayes (alpha = 1.0) classifier was presented for the primary analyses (43). Two additional classifiers were trained and tested given their ability to fit increasingly complex relationships between the features: Support Vector Machine (RBF kernel, L2-penalty, C = 0.5), and an Artificial Neural Network (ANN) with one hidden layer (256 hidden units), ReLU activation, and an L2-penalty of 5*10^−4^. The ANN was trained using an Adam optimizer with initial learning rate 1*10^−3^, batch size 64 (shuffled on each iteration), and early stopping after 20 iterations with no loss improvement on a held-out 10% validation set.

The train-test split for each classifier was established using a leave-one-subject-out-cross-validation (LOSOCV) approach, such that each classifier was trained on the data from all subjects except for one and was used to predict the trials of that left-out subject. This was performed across all subject-folds, and classifier predictions were aggregated across participants. Confusion matrices were calculated using this aggregated participant data, and accuracy was calculated as the ratio of the trace of the confusion matrix to the sum of all its entries. Accuracy was evaluated for statistical significance using a non-parametric permutation test. The above process was repeated (n=10,000) using shuffled target labels in order to establish a null distribution, and test accuracy was assessed for significance against this distribution.

### Data Visualization and Statistical Analysis

Template cortical surface visualization was performed via FieldTrip (44) (https://github.com/fieldtrip/fieldtrip) and lesion overlap maps were generated using svrlsmgui (45) (https://github.com/atdemarco/svrlsmgui). Linear mixed effects modeling was used to perform statistical comparisons with repeated measures via the nlme package in R (https://cran.r-project.org/web/packages/nlme/citation.html). The signal’s origin (i.e., normal-appearing/glioma-infiltrated cortex) was modeled as a fixed effect and the participants were modeled as random effects. Significance between contrast estimates were calculated using the multcomp package in R (https://cran.r-project.org/web/packages/multcomp/index.html). For continuous variables without repeated measures, t-tests were used. A threshold of p < 0.05 was used to denote statistical significance and corrections for multiple comparisons were made using the Bonferroni method.

## Acknowledgments and Funding Sources

This study was supported by the NIH grants K08NS110919 and P50CA097257, the Robert Wood Johnson Foundation grant 74259, the UCSF LoGlio Collective, and the Sheri Sobrato Brisson Brain Cancer Fund to S.H-J.; Sullivan Brain Cancer Fund to S.K.; the NIH grant R00DC013828 to D.B.

## Supplementary Information for

**Figure S1:**
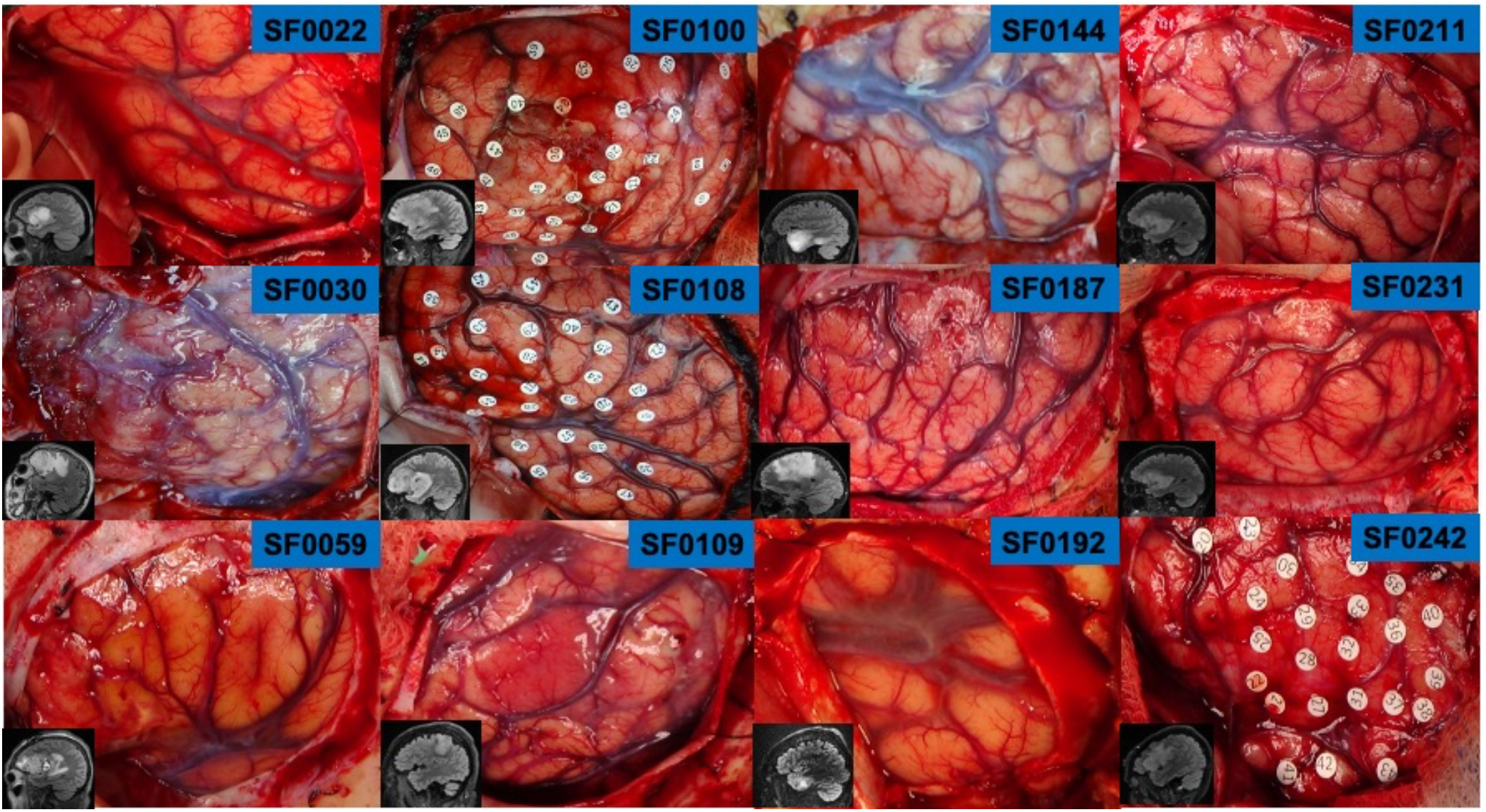
Intraoperative photographs of the cortical regions exposed for each participant. Insets demonstrate T2-weighted fluid attenuated inversion recovery (FLAIR) sequences with regions of signal hyperintensity representing glioma-infiltrated cortex. Gliomas project to the cortical surface in all twelve participants, facilitating electrocorticographic sampling from both glioma-infiltrated and normal-appearing cortex.

**Figure S2:**
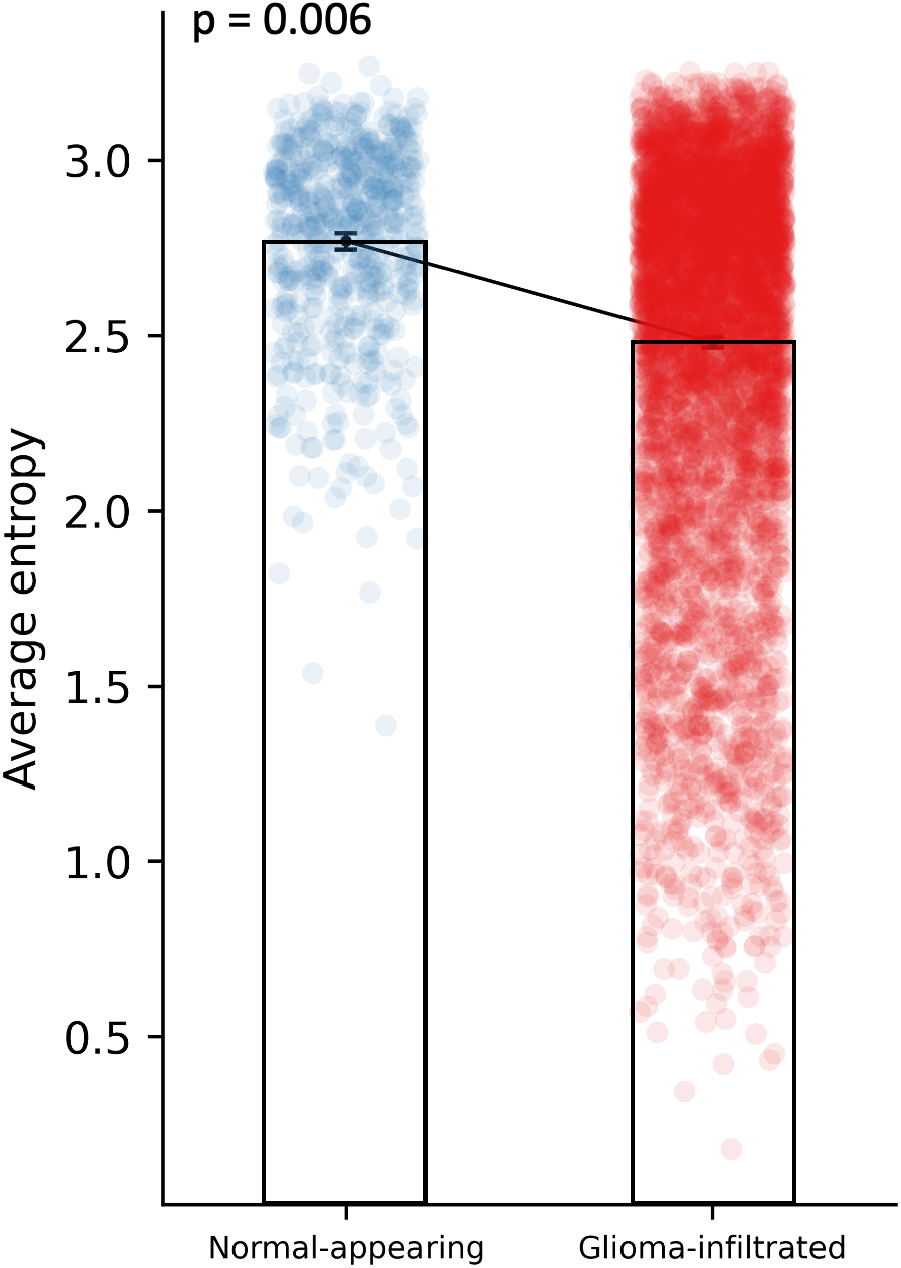
Difference in average trial entropy between normal-appearing and glioma-infiltrated cortex using Shannon’s metric for information. Statistical testing was performed using a linear mixed-effects model in which the type of cortex (i.e., normal-appearing, glioma-infiltrated) was modeled as fixed effects and participant labels were modeled as random effects. Neural signals from glioma-infiltrated cortex have lower entropy on average and therefore less capacity for information encoding.

**Figure S3:**
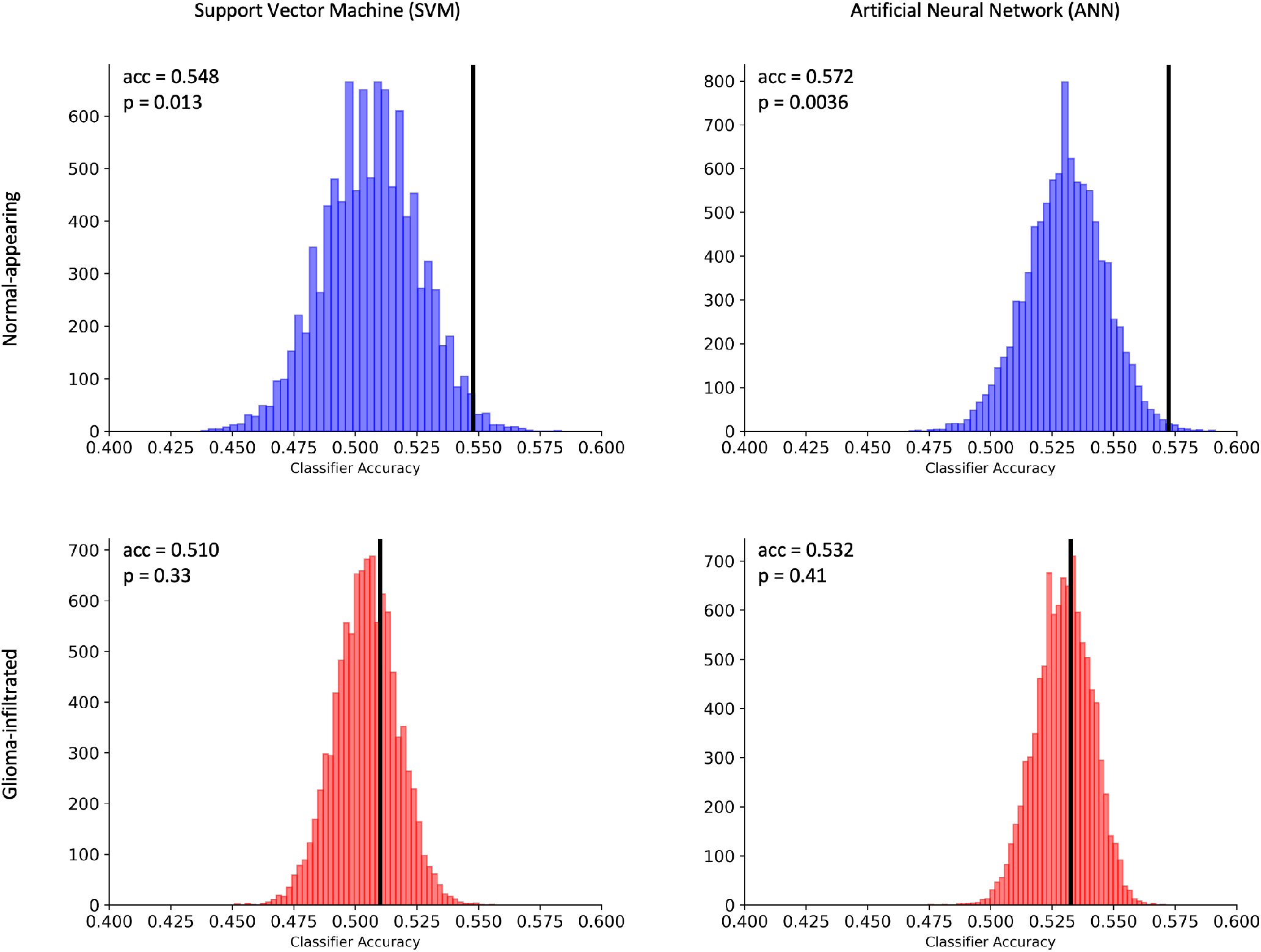
Neural decoding results using signals from normal-appearing (top) and glioma-infiltrated (bottom) cortex with two different classifiers: support vector machine (SVM, left) and artificial neural network (ANN, right). Actual model performance is demonstrated by the black bar in each plot. Histograms demonstrate the results of 10,000 permutations of randomly shuffled labels corresponding to monosyllabic and polysyllabic trial conditions and thereby the null distributions for nonparametric testing. Significance was determined by comparing this performance against the respective null distribution. Above-chance decoding was achieved using both SVM and ANN in normal-appearing cortex, but not in glioma-infiltrated cortex.

